# A comparative analysis of SARS-CoV-2 antivirals in human airway models characterizes 3CL^pro^ inhibitor PF-00835231 as a potential new treatment for COVID-19

**DOI:** 10.1101/2020.08.28.272880

**Authors:** Maren de Vries, Adil S Mohamed, Rachel A Prescott, Ana M Valero-Jimenez, Ludovic Desvignes, Rebecca O’Connor, Claire Steppan, Joseph C Devlin, Ellie Ivanova, Alberto Herrera, Austin Schinlever, Paige Loose, Kelly Ruggles, Sergei B Koralov, Annaliesa S. Anderson, Joseph Binder, Meike Dittmann

## Abstract

Severe acute respiratory syndrome coronavirus 2 (SARS-CoV-2) is the etiological agent of Coronavirus Disease 2019 (COVID-19). There is a dire need for novel effective antivirals to treat COVID-19, as the only approved direct-acting antiviral to date is remdesivir, targeting the viral polymerase complex. A potential alternate target in the viral life cycle is the main SARS-CoV-2 protease 3CL^pro^ (M^pro^). The drug candidate PF-00835231 is the active compound of the first anti-3CL^pro^ regimen in clinical trials. Here, we perform a comparative analysis of PF-00835231, the pre-clinical 3CL^pro^ inhibitor GC-376, and the polymerase inhibitor remdesivir, in alveolar basal epithelial cells modified to express ACE2 (A549^+ACE2^ cells). We find PF-00835231 with at least similar or higher potency than remdesivir or GC-376. A time-of-drug-addition approach delineates the timing of early SARS-CoV-2 life cycle steps in A549^+ACE2^ cells and validates PF-00835231’s early time of action. In a model of the human polarized airway epithelium, both PF-00835231 and remdesivir potently inhibit SARS-CoV-2 at low micromolar concentrations. Finally, we show that the efflux transporter P-glycoprotein, which was previously suggested to diminish PF-00835231’s efficacy based on experiments in monkey kidney Vero E6 cells, does not negatively impact PF-00835231 efficacy in either A549^+ACE2^ cells or human polarized airway epithelial cultures. Thus, our study provides *in vitro* evidence for the potential of PF-00835231 as an effective SARS-CoV-2 antiviral and addresses concerns that emerged based on prior studies in non-human *in vitro* models.

**Importance:** The arsenal of SARS-CoV-2 specific antiviral drugs is extremely limited. Only one direct-acting antiviral drug is currently approved, the viral polymerase inhibitor remdesivir, and it has limited efficacy. Thus, there is a substantial need to develop additional antiviral compounds with minimal side effects and alternate viral targets. One such alternate target is its main protease, 3CL^pro^ (M^pro^), an essential component of the SARS-CoV-2 life cycle processing the viral polyprotein into the components of the viral polymerase complex. In this study, we characterize a novel antiviral drug, PF-00835231, which is the active component of the first-in-class 3CL^pro^-targeting regimen in clinical trials. Using 3D *in vitro* models of the human airway epithelium, we demonstrate the antiviral potential of PF-00835231 for inhibition of SARS-CoV-2.

## Introduction

In December 2019, multiple cases of severe pneumonia with unexplained etiology were reported in Wuhan, China^1^. The infectious agent was identified as a novel member of the family *Coronaviridae*^1^, later named severe acute respiratory syndrome coronavirus 2 (SARS-CoV-2)^2^. The resulting disease, Coronavirus Disease 2019 (COVID-19), has since become a deadly pandemic.

A number of candidate drugs that may inhibit SARS-CoV-2 infection and replication have been proposed. However, only one direct-acting antiviral is currently approved for the treatment of COVID-19: remdesivir, a nucleoside analog that inhibits the SARS-CoV-2 RNA-dependent RNA-polymerase (RdRp). Remdesivir is incorporated into viral RNA by the RdRp, resulting in chain termination of both viral transcripts and *de novo* synthesized viral genomes^3^. Given this considerably limited arsenal of direct-acting antivirals for COVID-19, it remains a strategic priority to develop novel compounds with minimal side effects and that are directed against alternate viral targets.

One such alternate SARS-CoV-2 target is its main protease, 3CL^pro^ (M^pro^), which plays an essential role in the viral life cycle: Upon entry and uncoating of the viral particles, the positive-stranded RNA genome is rapidly translated into two polyproteins which are subsequently processed into functional proteins by PL2^pro^ and 3CL^pro^ viral proteases^4^. 3CL^pro^ is the main protease and is responsible for releasing 11 of the 13 individual proteins, including the polymerase subunits, enabling their proper folding and assembly into the active polymerase complex^5^. Thus, blocking 3CL^pro^ activity effectively shuts down the life cycle before viral transcription or replication occur, making it an enticing target for intervention^6^. In addition, 3CL^pro^ has a unique substrate preference (Leu-Gln ↓ {Ser, Ala, Gly}), a preference not shared by any known human protease, implying the potential for high selectivity and low side effects of 3CL^pro^-targeting drugs^7^. Although there have been intense efforts to develop 3CL^pro^ inhibitors specific for SARS-CoV-2^6–13^, only one inhibitor has been brought to the clinic, PF-07304814, which is the first anti-3CL^pro^ compound in clinical trials.

PF-07304814 is a ketone-based covalent cysteine protease inhibitor^9^. It is administered as a phosphate prodrug, which is then metabolized to its active form, PF-00835231^14^. PF-00835231 was initially designed in response to a previous coronavirus epidemic in 2003, as an inhibitor for the 3CL^pro^ of SARS-CoV^9^. However, with SARS-CoV disease declining, clinical studies were not practical and, consequently, PF-00835231 was never tested in patients. Because 3CL^pro^ of SARS-CoV and SARS-CoV-2 are 96% identical at the amino acid level, including 100% identity within the catalytic pocket^7^, it seemed reasonable to assume that PF-00835231 may inhibit SARS-CoV-2 as well.

Indeed, a recent study demonstrated antiviral activity of PF-00835231 against SARS-CoV-2, albeit at high micromolar levels^14^. The study was performed in Vero E6 cells, a monkey kidney cell line in which SARS-CoV-2 replicates to high titers, but which is known to express high levels of the efflux transporter P-glycoprotein (also known as Multi-Drug Resistance Protein 1, MDR1, and encoded by gene ATP Binding Cassette Subfamily B Member 1, *ABCB1*)^15^. Inhibiting MDR1 function significantly increased antiviral efficacy in Vero E6 cells, suggesting that PF-00835231 is an MDR1 substrate^14^. MDR1 is well-studied in the context of human immunodeficiency virus 1 (HIV-1) protease inhibitors such as lopinavir or ritonavir, where it reduces intracellular protease inhibitor levels and contributes to drug resistance in T-cells and monocytes^16^. In contrast to HIV-1, where viral replication is essentially limited to T-cells and monocytes, SARS-CoV-2 infects multiple organs and cell types throughout the human body, with the first and major site of replication being cells of the respiratory tract^17, 18^. Thus, to investigate the potential role of MDR1 on antiviral potency of PF-00835231 against SARS-CoV-2, it is imperative to utilize experimental model systems representing the human airways.

Here, we characterize the antiviral potency and cytotoxicity profile of PF-00835231 in two human airway models: a human type II alveolar epithelial cell line, and polarized human airway epithelial cultures. In side-by-side experiments, we place PF-00835231’s antiviral efficacy against SARS-CoV-2 in context of another, pre-clinical 3CL^pro^ inhibitor, GC-376, and of the current standard-of-care, remdesivir. Finally, we address the impact of P-glycoprotein (MDR1) on PF-00835231’s antiviral efficacy in the human airways.

## Results

### Establishing A549^+ACE2^ cells as a tool to determine SARS-CoV-2 infection and cytopathic effect by high-content microscopy

The human adenocarcinoma alveolar epithelial cell line A549 is a workhorse cell line in the study of respiratory viruses. However, A549 cells are not permissive to SARS-CoV-2 infection, as they do not highly express the SARS-CoV-2 receptor ACE2^19^. To make A549 cells amenable for experiments with SARS-CoV-2, we generated a stable A549 cell line expressing ACE2 exogenously. We confirmed elevated levels of ACE2 mRNA in A549^+ACE2^ cells by RT-qPCR, and of ACE2 protein by Western blot, flow cytometry, and confocal microscopy (Fig. 1a-e).

**Figure 1.**
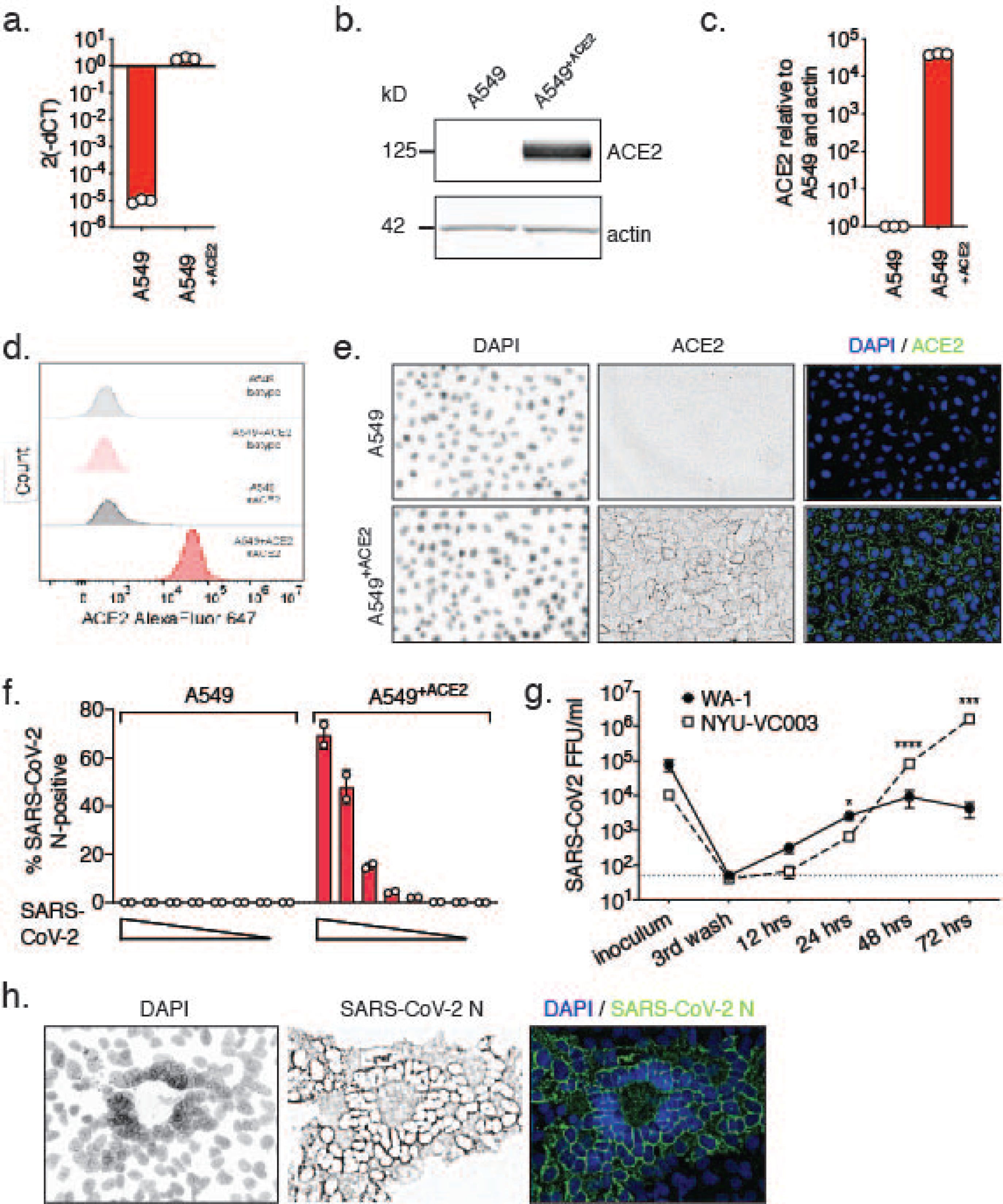
Validation of A549^+ACE2^ cells as a tool to study SARS-CoV-2. A549^+ACE2^ cells were generated by lentiviral transduction delivering an ACE2 overexpression construct and subsequent bulk-selection. **a.-e.** ACE2 expression in A549 parental or A549^+ACE2^ cells determined by RT-qPCR (**a.**), western blot (**b.,** quantified in **c.**), flow cytometry (**d.**), or microscopy (**e.**). **f.** A549 parental or A549^+ACE2^ cells were infected with a serial dilution of SARS-CoV-2 USA-WA1/2020. At 24 h, cells were fixed, stained for SARS-CoV-2 N protein, and infected cells were quantified by high-content microscopy. Means ± SEM from duplicate wells. **g.** A549^+ACE2^ cells were infected with SARS-CoV-2 USA-WA1/2020 or USA/NYU-VC-003/2020 at MOI 0.01, and infectious progeny titers, collected from supernatants over time, determined by focus forming assay on A549^+ACE2^ cells. Means ± SEM from n=3 independent experiments. Unpaired t-test, *p<0.05, ***p<0.001, ****p<0.0001. **h.** Confocal microscopy of SARS-CoV-2 syncytia in A549^+ACE2^ cells at 48 hpi.

To determine permissiveness, we infected A549 or A549^+ACE2^ cells with a serial dilution of SARS-CoV-2, in a 96-well format, for 24 h. Using immunofluorescence staining for SARS-CoV-2 nucleocapsid protein (N) and high-content microscopy, we found A549^+ACE2^ cells permissive to SARS-CoV-2 infection, whereas parent A549 cells were not (Fig. 1f).

Since the discovery of SARS-CoV-2, limited evolution has been observed, which has been attributed to the proof-reading mechanism of coronavirus polymerases^20^. The two major lineages of SARS-CoV-2 circulating globally as of time of writing are represented by the Wuhan basal clade and the spike protein D614G clade, also referred to as clades A and B, respectively^21^. Compared to clade A, clade B isolates carry a mutation in the spike-encoding gene S, which results in amino acid substitution D614G. D614G is frequently accompanied by an additional mutation in ORF 1b, which encodes the RNA-dependent RNA-polymerase complex (RdRp), resulting in substitution P323L in the polymerase subunit NSP12^22^. Clade B viruses are more prevalent globally, which might be due to their increased efficiency infecting cells in the upper respiratory tract and subsequently increased transmissibility, enabled by the Spike D614G mutation^23, 24^.

To characterize viral growth of representatives from the two major clades in our model, we challenged A549^+ACE2^ cells with the clinical SARS-CoV-2 isolate USA-WA1/2020, a clade A representative^25^, or with USA/NYU-VC-003/2020, a clade B representative, the latter of which we had isolated in March 2020^26^. USA/NYU-VC-003/2020 carries both of the signature clade B amino acid changes, S D614G and NSP12 P323L, but its 3CL^pro^ sequence is identical to that of USA-WA1/2020. In low MOI growth kinetics on A549^+ACE2^ cells, we found that growth of clade B USA/NYU-VC-003/2020 exceeds that of clade A USA-WA1/2020, especially at later times of infection (Fig. 1g). We were able to detect *de novo* produced infectious particles as soon as 12 hours post infection (hpi) for USA-WA1/2020, suggesting that the SARS-CoV-2 life cycle in A549^+ACE2^ cells is completed by that time. In terms of producing infectious titers, USA/NYU-VC-003/2020 initially lagged behind USA-WA1/2020, but then yielded significantly higher titers at 48 and 72 hpi.

Finally, we observed that the cytopathic effect (CPE) caused by SARS-CoV-2 on A549^+ACE2^ cells manifests in syncytia formation, in which the nuclei form a ring-like structure (Fig. 1h). This effect had previously been described for other coronaviruses^27, 28^, although the exact mechanism for the ring-like nuclear structure formation remains to be elucidated. Altogether, our data establish A549^+ACE2^ cells as a tractable tool to study SARS-CoV-2 infection, spread, and cytopathic effect.

### PF-00835231 potently inhibits SARS-CoV-2 in A549^+ACE2^ cells

PF-00835231 is the active compound of the first anti 3CL^pro^ regimen currently tested in clinical trials^9^. We studied and compared three compounds in regards to SARS-CoV-2 antiviral activity and cytotoxicity: i. PF-00835231, ii. the pre-clinical 3CL^pro^ inhibitor GC-376, which is licensed for veterinary use in Feline Coronavirus infections^29^ and recently shown to inhibit SARS-CoV-2 in Vero E6 cells^26^, and iii. the polymerase inhibitor remdesivir, which is currently the only direct-acting antiviral approved in the US to treat SARS-CoV-2 infections, and is thus standard-of-care. We exposed A549^+ACE2^ cells with escalating doses of the three respective drugs, challenged them with SARS-CoV-2, and measured virus antigen (N)-expressing cells by high-content microscopy (Fig. 2a). In parallel, we determined cellular viability by measuring ATP levels in drug-treated, but uninfected cells. Antiviral assays were performed with both our clade A and clade B SARS-CoV-2 representatives.

**Figure 2.**
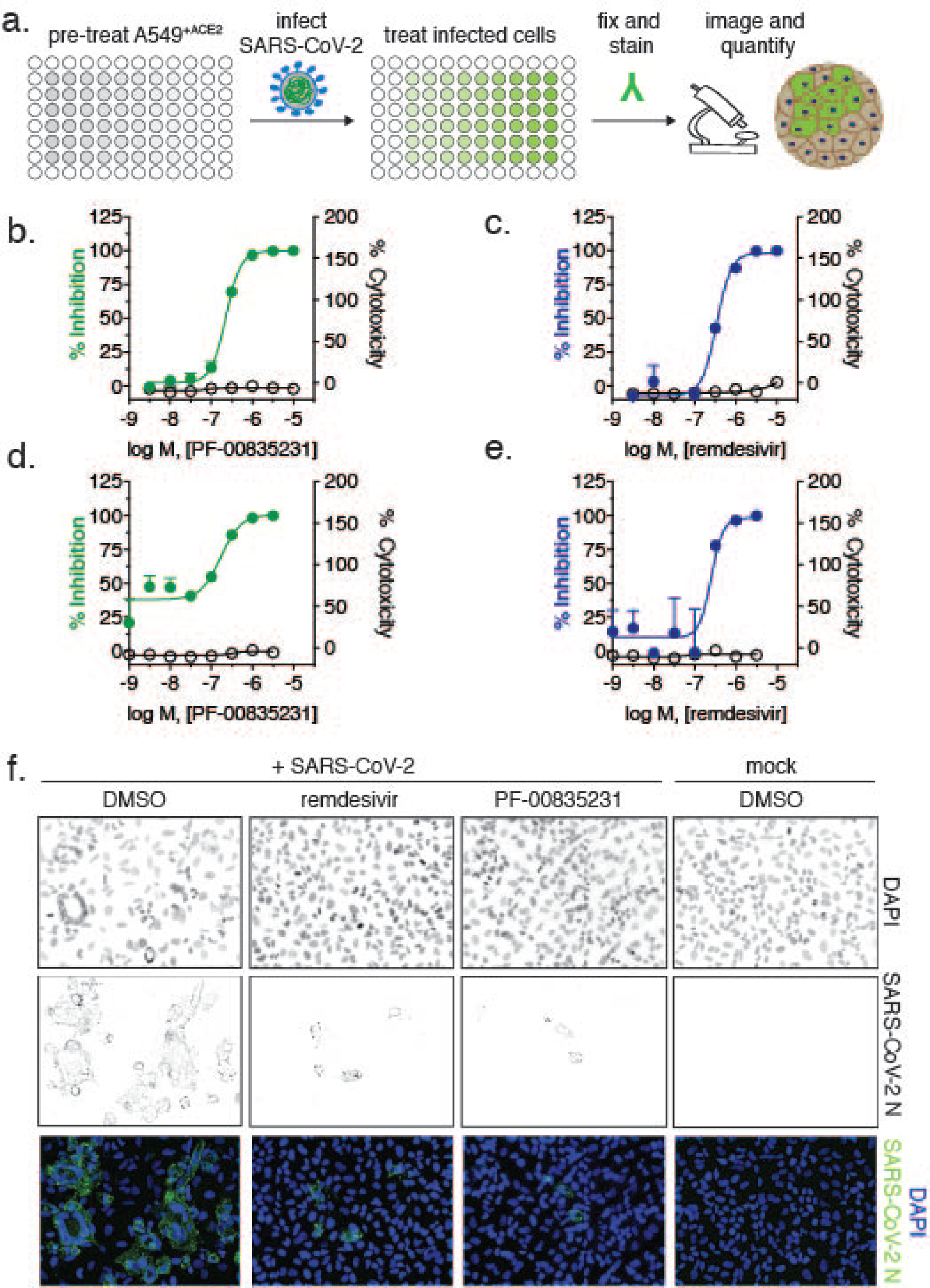
Antiviral SARS-CoV-2 activity and cytotoxicity of PF-00835231, and remdesivir in A549^+ACE2^ cells. **a.** Antiviral assay workflow. A549^+ACE2^ cells were pretreated with serial dilutions of PF-00835231 or remdesivir, then infected with SARS-CoV-2 while continuing drug treatment. At 24 or 48 h, cells were fixed, stained for SARS-CoV-2 N protein, and infected cells quantified by high-content microscopy. Cytotoxicity was measured in similarly treated but uninfected cultures via CellTiter-Glo assay. EC_50_, EC_90_ and CC_50_ from n=3 independent experiments are listed in Table 1. **b.** PF-00835231, and **c.** remdesivir antiviral activity and cytotoxicity in A549^+ACE2^ cells infected with SARS-CoV-2 USA-WA1/2020 for 24 h. Representative graphs shown. **d.** PF-00835231, and **e.** remdesivir antiviral activity and cytotoxicity in A549^+ACE2^ cells infected with SARS-CoV-2 USA/NYU-VC-003/2020 for 24 h. Representative graphs shown. **f.** Representative images of SARS-CoV-2 USA-WA1/2020 syncytia formation at 48 hpi in A549^+ACE2^ cells under treatment with 0.33 µM PF-00835231, or remdesivir.

**Table 1.**
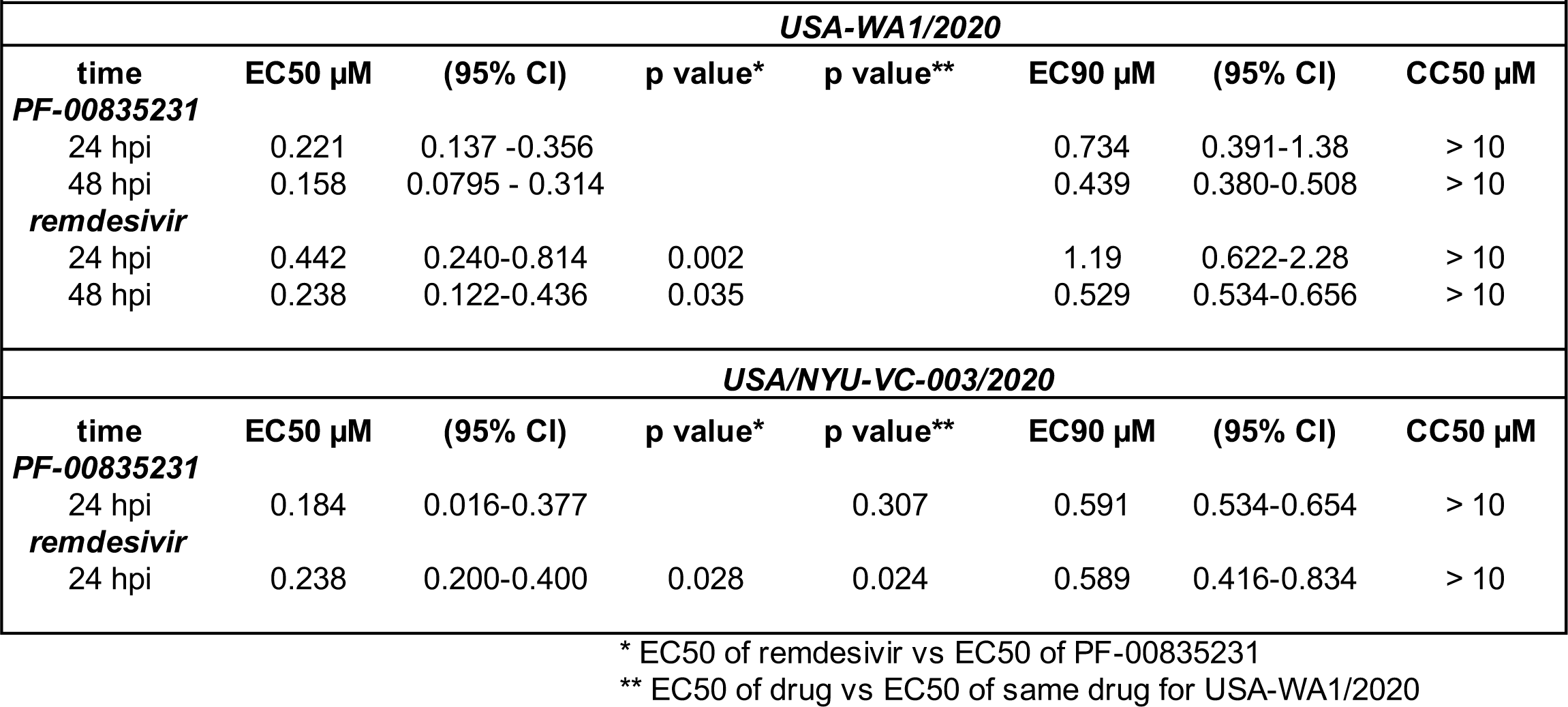
Antiviral efficacy and cytotoxicity of PF-00835231 versus remdesivir on A549+ACE2 cells

In a first set of experiments, we compared antiviral efficacy between PF-00835231 and remdesivir side-by-side (Fig. 2 and Tab.1). PF-00835231 inhibited the clade A representative SARS-CoV-2 USA-WA1/2020 with an average 50% effective concentration (EC_50_) of 0.221 µM at 24 h, and 0.158 µM at 48 h (Fig. 2b, Tab. 1). As such, PF-00835231 was statistically more potent than remdesivir with an EC_50_ 0.442 µM at 24 h, and 0.238 µM at 48 h (Fig. 2c, Tab. 1). None of the compounds showed detectable cytotoxicity (Fig. 2b-e, Tab. 1). We then compared antiviral efficacy of PF-00835231 and remdesivir for clade B USA/NYU-VC-003/2020. Due to cytopathic effects driven by USA/NYU-VC-003/2020 at the 48 h timepoint, we only determined antiviral efficacy at 24 h. PF-00835231 was inhibitory with an EC_50_ of 0.184 µM, and was thus again statistically more potent than remdesivir with EC_50_ of 0.283 µM.

Interestingly, only the polymerase inhibitor remdesivir exhibited statistically significantly weaker antiviral activity against the clade A isolate compared to the clade B isolate, with EC_50_ of 0.442 µM (vs clade A) and EC_50_ of 0.238 µM (vs clade B; Tab. 1). This might be an impact of the polymerase subunit NSP12 P323L mutation present in the clade B representative. Next, we analyzed microscopy data for drug-mediated inhibition of the CPE, including ring-shaped syncytia formation. PF-00835231 and remdesivir both decreased the overall number of infected foci, and fully protected A549^+ACE2^ cells from ring syncytia formation, at 0.33 µM and above (Fig. 2f and not shown).

In a second set of experiments, we compared antiviral efficacy between PF-00835231 and GC-376 side-by-side (Fig. 3a-d and Tab. 2). PF-00835231 inhibited the clade A representative SARS-CoV-2 USA-WA1/2020 with an EC_50_ of 0.422 µM at 24 h, and 0.344 µM at 48 h (Fig. 2a, Tab. 1). This slight shift in EC_50_ values as compared to those in Table 1 can be explained by intra-assay variation, which is higher for live cell assays as it is for binding assays. For this reason we could not compare PF-00835231 across assays. In the direct comparison, PF-00835231 trended towards being more potent than GC-376, which exhibited an EC_50_ of 0.632 µM at 24 h, and 0.696 µM at 48 h (Fig. 3b, Tab. 2). For clade B USA/NYU-VC-003/2020, PF-00835231 was inhibitory with an EC_50_ of 0.326 µM, and thus again trended towards being more potent than GC-376 with an EC_50_ of 0.529 µM (Fig. 3c-d, Tab. 2).

**Figure 3.**
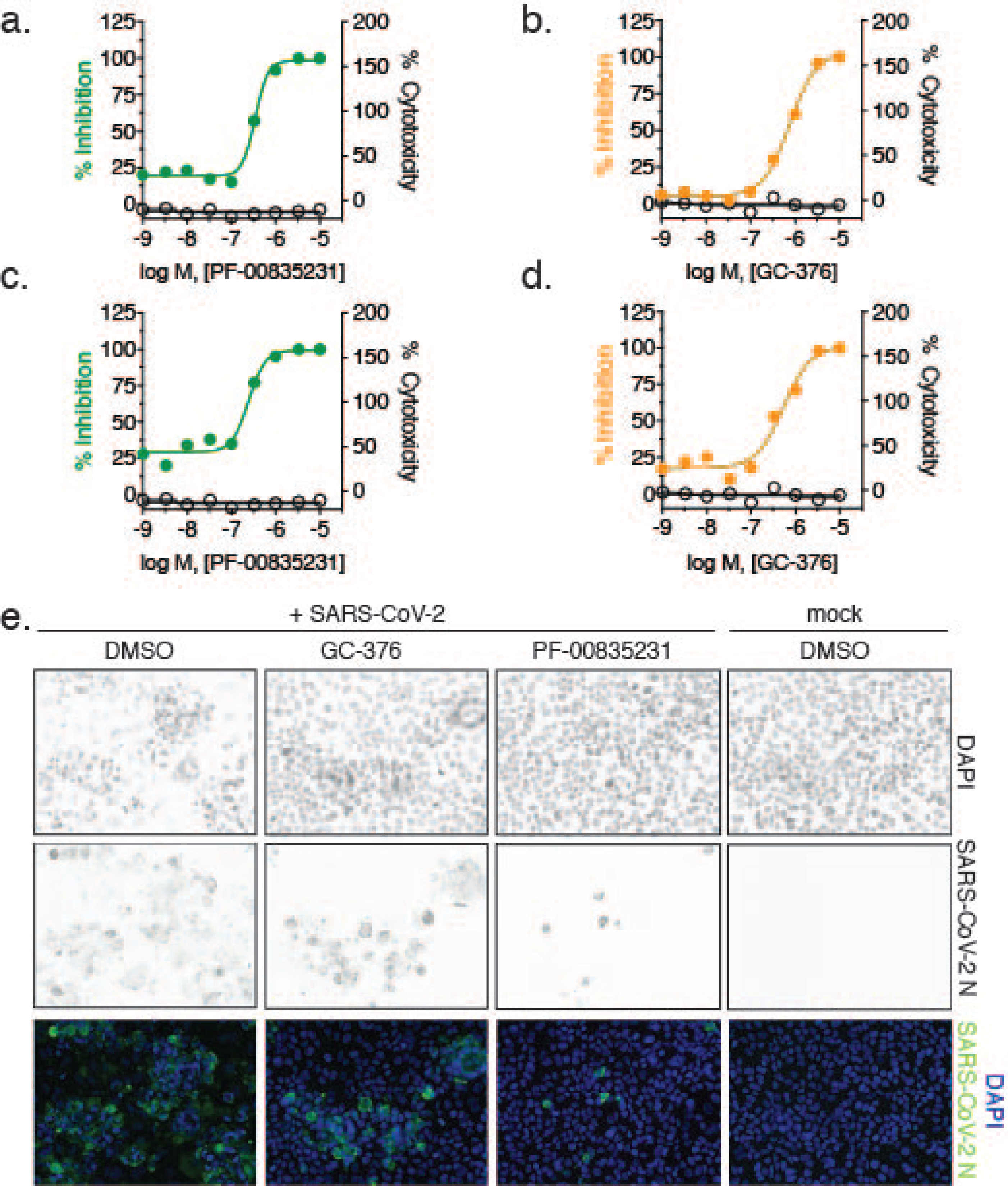
Antiviral SARS-CoV-2 activity and cytotoxicity of PF-00835231, and GC-376 in A549^+ACE2^ cells. Antiviral assay workflow as described in Fig. 2a. **a.** PF-00835231, and **b.** GC-376 antiviral activity and cytotoxicity in A549^+ACE2^ cells infected with SARS-CoV-2 USA-WA1/2020 for 24 h. Representative graphs shown. **c.** PF-00835231, and **d.** GC-376 antiviral activity and cytotoxicity in A549^+ACE2^ cells infected with SARS-CoV-2 USA/NYU-VC-003/2020 for 24 h. Representative graphs shown. **e.** Representative images of SARS-CoV-2 USA-WA1/2020 syncytia formation at 48 hpi in A549^+ACE2^ cells under treatment with 1 µM PF-00835231, or GC-376.

**Table 2.** Antiviral efficacy and cytotoxicity of PF-00835231 versus GC-376 on A549+ACE2 cells

|  |  |  |  |  |  |  |  |
| --- | --- | --- | --- | --- | --- | --- | --- |
| *USA-WA1/2020* |  |  |  |  |  |  |  |
| time | EC50 μM | (95% CI) | p value* | p value** | EC90 μM | (95% CI) | CC50 μM |
| **PF-00835231** |  |  |  |  |  |  |  |
| 24 hpi | 0.422 | 0.0836 - 2.13 |  |  | 0.978 | 0.326 - 2.93 | >10 |
| 48 hpi | 0.344 | 0.0842 - 1.404 |  |  | 1.158 | 0.358 - 3.74 | >10 |
| **GC-376** |  |  |  |  |  |  |  |
| 24 hpi | 0.623 | 0.257 - 1.506 | 0.366 |  | 4.55 | 1.89 - 10.91 | >10 |
| 48 hpi | 0.696 | 0.198 - 2.44 | 0.114 |  | 5.25 | 2.53 - 10.875 | >10 |
| *USA/NYU-VC-003/2020* |  |  |  |  |  |  |  |
| time | EC50 μM | (95% CI) | p value* | p value** | EC90 μM | (95% CI) | CC50 μM |
| **PF-00835231** |  |  |  |  |  |  |  |
| 24 hpi | 0.326 | 0.098 - 1.08 |  | 0.543 | 1.17 | 0.115 - 11.89 | >10 |
| **GC-376** |  |  |  |  |  |  |  |
| 24 hpi | 0.529 | 0.184 - 1.512 | 0.265 | 0.700 | 2.734 | 0.897 - 8.33 | >10 |

\* EC50 of GC-376 vs EC50 of PF-00835231
\*\* EC50 of drug vs EC50 of same drug for USA-WA1/2020

Both protease inhibitors PF-00835231 and GC-376 had similar antiviral activities between the two clades in this assay (Tab. 1 and Tab. 2). This is in line with the fact that 3CL^pro^ is identical in these two viruses. Finally, GC-376 decreased the number and size of viral foci, but was unable to protect cells from virus-induced CPE at 1 µM (Fig. 3e). In contrast, at 1 µM and above, PF-00835231 fully protected A549^+ACE2^ cells from CPE (Fig. 3e and not shown).

Collectively, we show that, in this assay, PF-00835231 inhibits isolates from both major SARS-CoV-2 lineages at similar or better effective concentrations than remdesivir and the pre-clinical 3CL^pro^ inhibitor GC-376.

### Timing of PF-00835231 antiviral action against USA-WA1/2020 in A549^+ACE2^ cells is consistent with PF-00835231’s role as a 3CL^pro^ inhibitor

PF-00835231 and remdesivir target different SARS-CoV-2 proteins^9, 31^. PF-00835231 targets 3CL^pro^, blocking polyprotein processing and thus formation of the viral polymerase complex^32^. Remdesivir acts on the subsequent step, which is the incorporation of nucleotides into nascent viral RNA transcripts and genomes by the viral polymerase complex^3, 33^.

To determine whether the action of PF-00835231 is consistent with its established role as a 3CL^pro^ inhibitor, and to delineate the timing of early SARS-CoV-2 life cycle stages in A549^+ACE2^ cells, we performed time-of-drug-addition experiments^34^. This approach determines how long the addition of a drug can be delayed before it loses antiviral activity. Using one-hour-increments (from 1 h prior to 4 h post infection), we varied the time-of-drug-addition for a monoclonal neutralizing antibody (viral attachment inhibition control), GC-376 (3CL^pro^ inhibition control), PF-00835231, and remdesivir (RdRp inhibition control). We measured the percentage of SARS-CoV-2-infected cells via high-content microscopy at 12 hpi, which corresponds to one replication cycle in A549^+ACE2^ cells, as determined previously (Fig. 1g). We synchronized infection using a preincubation step at 4°C, followed by a transition to 37° C at 1 h post-addition of virus, and used the minimum treatment doses for each drug that led to undetectable infection levels – 3 µM for PF-00835231 and the neutralizing antibody, and 10 µM for remdesivir and GC-376.

The neutralizing antibody lost its antiviral function first, starting at the first addition point post-infection (1 h), confirming blockage of attachment and entry as the mode of antiviral action (Fig. 4a). Interestingly, all three drugs, GC-376, PF-00835231, and remdesivir lost antiviral action at the same time of addition, starting at 2 hpi, and with subsequently enhanced loss of activity at 3 and 4 hpi (Fig. 4a). This suggests that polyprotein processing and the start of viral transcription / translation follow each other very closely in time. These time-of-drug-addition experiments confirm the timing of PF-00835231’s antiviral action during the early stages of intracellular virus propagation, consistent with its role as a 3CL^pro^ inhibitor. Furthermore, these experiments delineate the timing of the SARS-CoV-2 life cycle events in the tissue culture model of A549^+ACE2^ cells (Fig. 4b) and demonstrate that polymerase and protease inhibitors such as PF-00835231 can effectively block SARS-CoV-2 replication in cells when administered within a few hours after infection.

**Figure 4.**
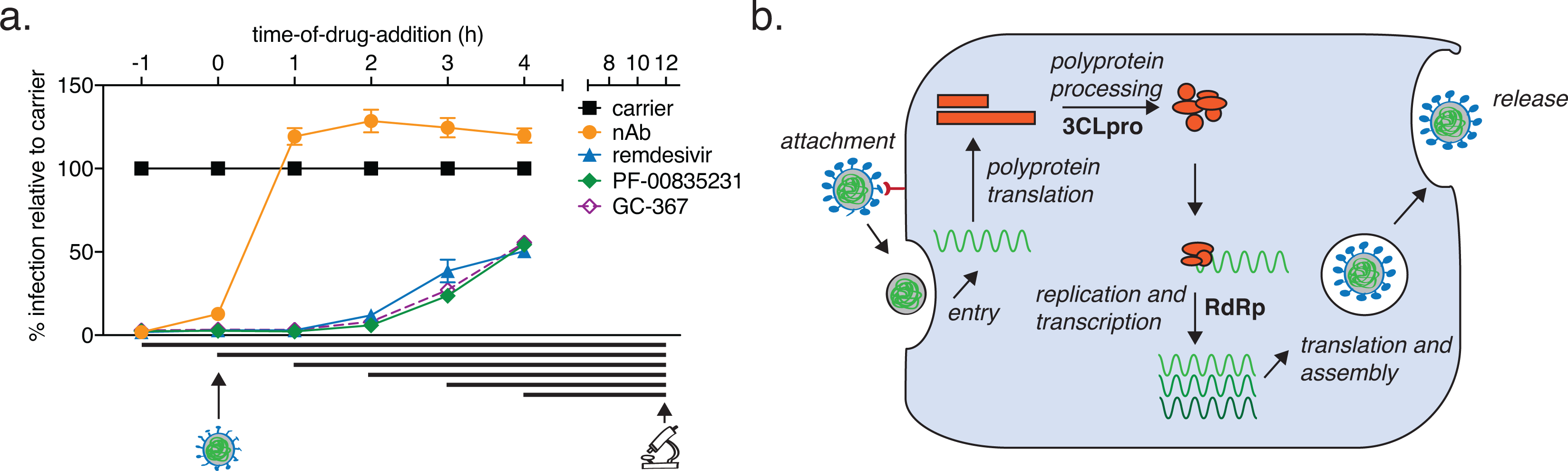
Time-of-drug-addition assay for PF-00835231, GC-376, remdesivir, and a neutralizing antibody in A549^+ACE2^ cells. **a.** At the indicated time points, A549^+ACE2^ cells were infected with SARS-CoV-2 USA-WA1/2020, treated with 3 µM monoclonal neutralizing antibody (control targeting attachment and entry), 10 µM of the drug GC-376 (control drug for 3CL^pro^ inhibition), 3 µM PF-00835231 (3CL^pro^ inhibitor), or 10 µM remdesivir (RdRp inhibitor). At 12 h (one round of replication) cells were fixed, stained for SARS-CoV-2 N protein, and infected cells quantified by high-content microscopy. Means ± SEM from n=3 independent experiments. **b.** Schematic of SARS-CoV-2 life cycle in A549^+ACE2^ cells. 3CL^pro^, 3C-like protease; RdRp, RNA-dependent RNA polymerase.

### PF-00835231 is well-tolerated in polarized human airway epithelial cultures (HAEC)

The human respiratory tract is a major entry portal for viruses, including SARS-CoV-2, and the first battle between host and virus occurs in cells of the respiratory epithelium. This specialized tissue contains three major cell types (basal, secretory, and ciliated) which are organized in a characteristic polarized architecture^35^. As shown, human adenocarcinoma alveolar epithelial A549^+ACE2^ cells are permissive for SARS-CoV-2 infection and allow for high-throughput experiments (Fig. 1-4). However, they do not recapitulate the complexity and architecture of the human airway epithelium.

To test PF-00835231 and remdesivir in an additional, more physiologically relevant, yet lower-throughput human model system, we generated polarized human airway epithelial cultures (HAEC). HAEC contain multiple cell types of the airway epithelium and recapitulate its typical architecture (Fig. 5a-d), which makes HAEC arguably one of the most physiologically relevant models for *in vitro* studies of human respiratory pathogens. HAEC are permissive to SARS-CoV-2 infections and were utilized to obtain the very first SARS-CoV-2 isolate in December 2019^1^.

**Figure 5.**
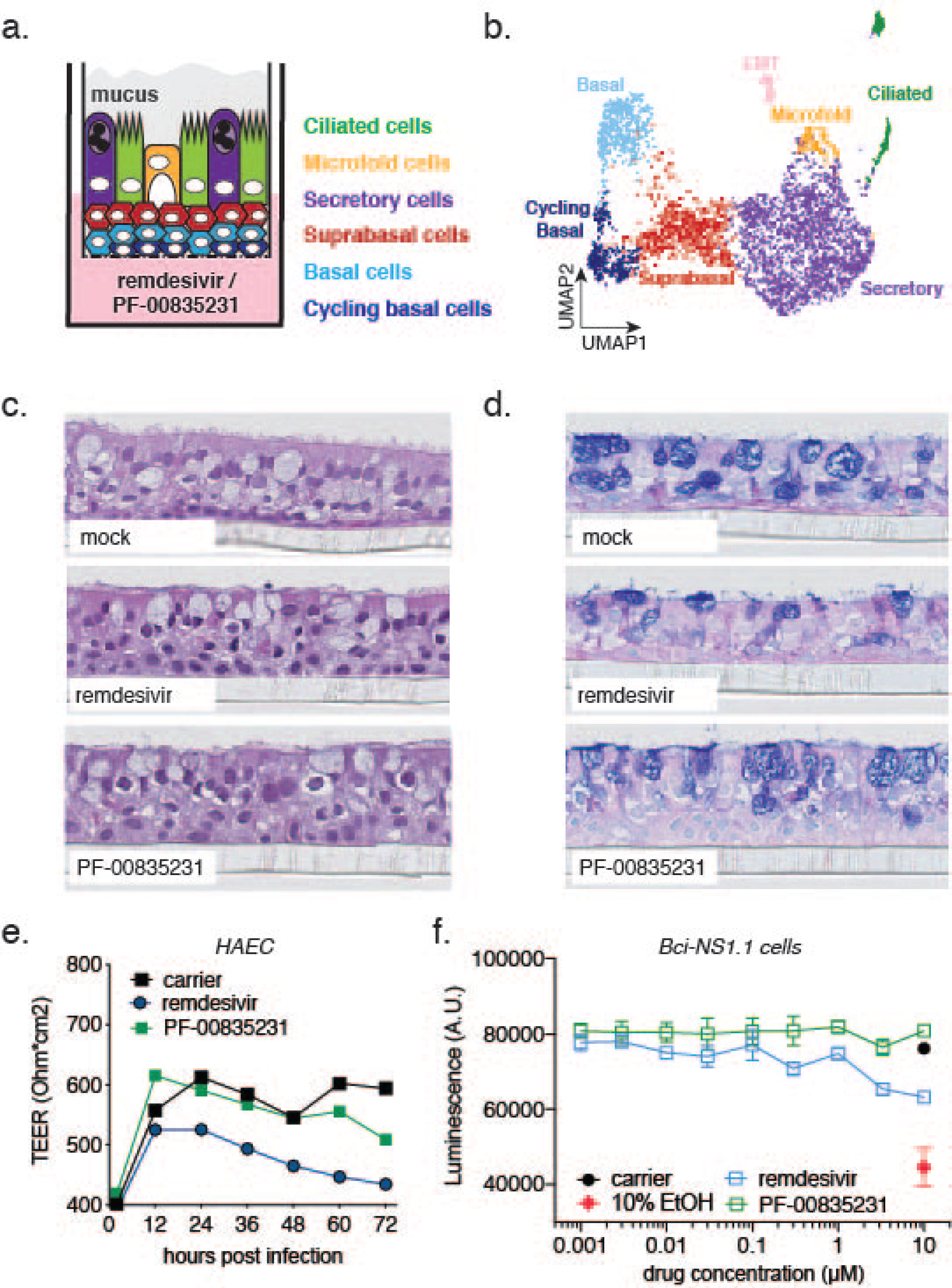
Cell composition of polarized human airway epithelial cultures (HAEC), and cytotoxicity of PF-00835231 and remdesivir. **a.** Schematic representation of a transwell containing a polarized HAEC in air-liquid interface. Dark blue, cycling basal cells; light blue, basal cells; red, suprabasal cells; purple, secretory cells; yellow, microfold cells; green, ciliated cells; grey, mucus. To test for cytotoxicity, drugs were added to the media in the basolateral chamber. **b.** Clustered UMAP of single cells determined by single-cell RNA sequencing from n=3 uninfected HAEC. Clusters were determined by markers from the literature^36, 37^ and by differentially expressed marker genes for each cluster determined by Wilcox test. **c., d.** Representative cross-sections of uninfected HAEC, 72 h post treatment with 10 µM PF-00835231 or 10 µM remdesivir. H&E (**c.**) or PAS-Alcian blue staining (**d.**). **e.** Trans-epithelial resistance (TEER) in drug-treated, uninfected HAEC over time as a measure of epithelial integrity. Means ± SEM from n=3 independent experiments. **f.** CellTiter-glo assay on undifferentiated, basal-like Bci-NS1.1 precursor cells. Means ± SEM from n=3 independent experiments.

First, we performed in-depth analyses of the cellular heterogeneity of our HAEC model system by single-cell RNA-sequencing (sc-RNAseq; Fig. 5b^38^). Gene-expression profiling enabled resolution of distinct clusters with cell types assigned based on previously published transcriptional signatures^36, 37^. We identified 7 different clusters as cycling basal, basal, suprabasal, secretory, ciliated, and microfold cells, as well as cells undergoing epithelial-mesenchymal transition (EMT, Fig. 5b). The EMT process in HAEC has previously been associated with loss of polarized organization as a consequence of remodeling^39^ and is likely to occur at a low level at steady-state in HAEC. Recapitulation of the major cell types and physiological conditions of the lung epithelium provided molecular confirmation for the HAEC system in assessing SARS-CoV-2 infection.

To establish the use of PF-00835231 in HAEC, we determined its cytotoxicity profile and compared it to that of remdesivir. We added PF-00835231 or remdesivir to the basolateral chamber of HAEC (Fig. 5a), and determined tissue morphology by histology and integrity of the epithelial layer by measuring trans-epithelial resistance (TEER; Fig. 5c-e). Neither drug caused measurable adverse effects on the morphology of the cultures (Fig. 5c,d). However, while remdesivir negatively impacted TEER over time, albeit not statistically significantly compared to untreated cultures, we did not observe this trend for PF-00835231 (Fig. 5e).

To complement our assessment of how well human epithelium tolerates these inhibitors, we took advantage of an alternative cytotoxicity assay on BCi-NS1.1 cells, the basal-like undifferentiated precursor cell monolayers used for generation of HAEC. We treated these monolayers with a dose range of PF-00835231 or remdesivir for 48 hours, and quantified ATP as a measure of cell viability, similar to previous experiments with A549^+ACE2^ cells. We did not detect a decrease in ATP upon PF-00835231 treatment, even at the highest amount of drug (10 µM) tested. In contrast, 10 µM of remdesivir caused a reduction in ATP levels compared to carrier control, albeit not statistically significantly (Fig. 5f). These experiments demonstrate that both drugs are well-tolerated in our model of polarized human airway epithelium.

### PF-00835231 exhibits potent anti-SARS-CoV-2 activity in HAEC

To determine PF-00835231’s anti-SARS-CoV-2 activity in HAEC, we added either 0.025, 0.5 or 10 µM PF-00835231 or remdesivir, or DMSO carrier control, to the basolateral chamber of HAEC (Fig. 6a-c). We then challenged HAEC apically with SARS-CoV-2 USA-WA1/2020, and determined viral infectious titers from apical washes collected at 12-hour increments.

**Figure 6.**
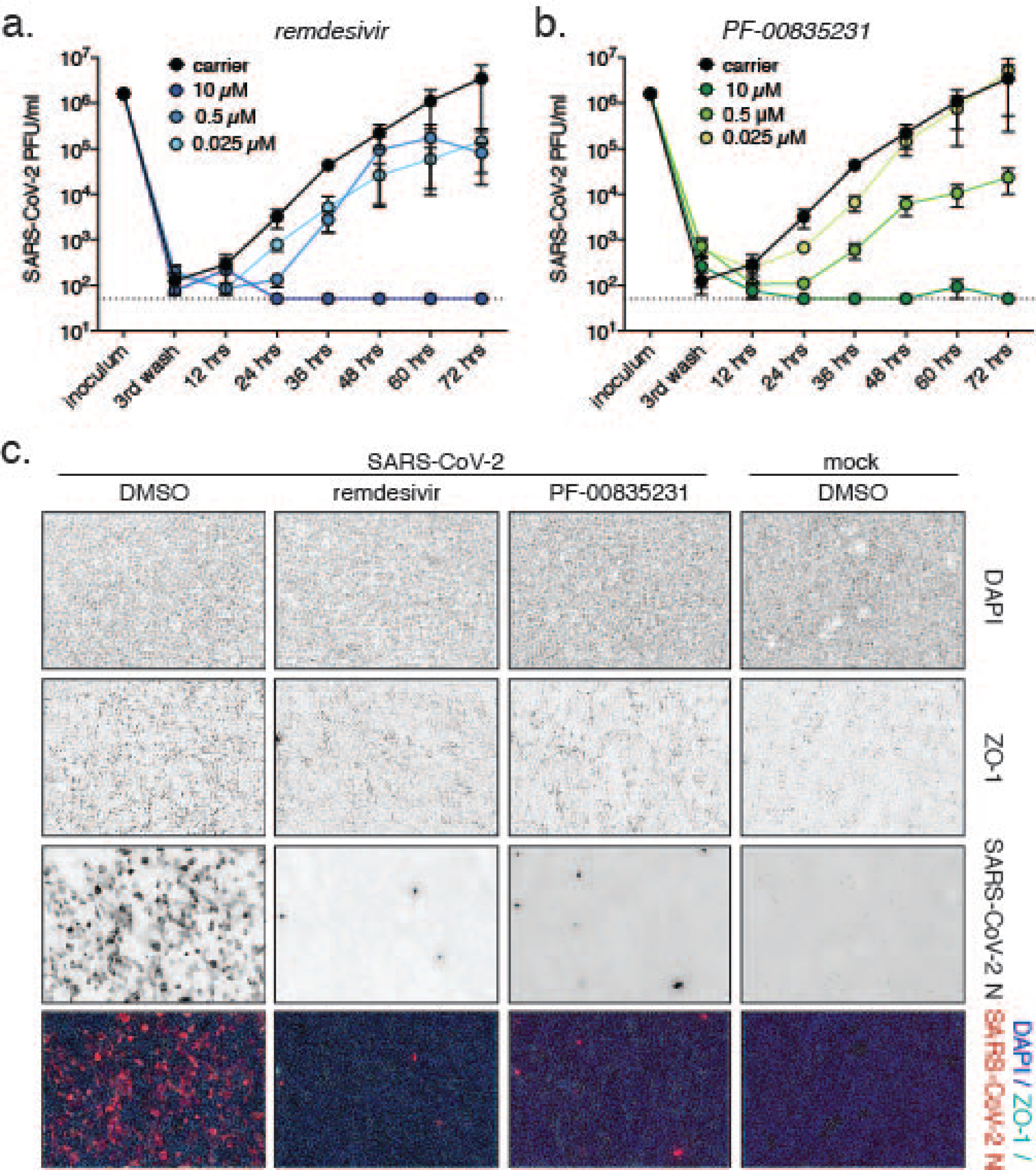
Comparative anti-SARS-CoV-2 activity of PF-00835231 and remdesivir in polarized human airway epithelial cultures (HAEC). To test for antiviral activity, drugs were added to the basolateral chamber, cultures infected with SARS-CoV-2 from the apical side, and apical washes collected in 12 h increments to determine viral titers by plaque assay. **a, b.** SARS-CoV-2 USA-WA1/2020 infectious titers from HAEC treated with incremental doses of remdesivir (**a.**) or PF-00835231 (**b.**). **c.** Representative top views of HAEC at 72 hpi; drug doses 0.3 µM. Blue, DAPI (nuclei); cyan, ZO-1 (tight junctions); red, SARS-CoV-2 N protein (infected).

We first detected progeny viral particles in apical washes from DMSO-treated cultures at 12 hpi (Fig. 6a, b), indicating that the SARS-CoV-2 life cycle in HAEC cells is completed by that time. Both PF-00835231 and remdesivir potently inhibited SARS-CoV-2 titers in a dose-dependent manner, with the 10 µM doses resulting in viral titers below the limit of detection at most time points (Fig. 6a, b).

To visualize SARS-CoV-2 infection in HAEC during drug treatment, we fixed infected HAEC at the 72 h endpoint and stained them for SARS-CoV-2-N-expressing cells (Fig. 6c). In carrier control cultures, we observed robust infection. Upon treatment with 10 µM PF-00835231 or remdesivir, we found in both cases the number of infected cells significantly reduced. Taken together, both remdesivir and PF-00835231 potently inhibit SARS-CoV-2 infection in our model of polarized human airway epithelium.

### Inhibiting the multi-drug transporter MDR1 does not increase efficacy of PF-00835231 in human airway epithelial cells

Previously, a hurdle in accurately determining PF-00835231’s *in vitro* efficacy was the action of the multi-drug efflux transporter P-glycoprotein (also known as MDR1 or *ABCB1*). However, these earlier studies were performed in the monkey kidney cell line Vero E6^9, 14^. MDR1 was found to efficiently export PF-00835231, thereby reducing intracellular PF-00835231 levels, and likely underestimating PF-00835231’s potency. In those studies, chemical inhibition of MDR1 in Vero E6 cells significantly increased PF-00835231’s antiviral efficacy^9, 14^.

Given the previously reported species differences in P-glycoprotein-mediated drug transport activity of MDR1 and the variability in expression levels of the *ABCB1* gene that encodes this drug transporter among cell types and tissues^40^, we sought to determine a potential role of MDR1 in our human *in vitro* airway models. We measured PF-00835231 anti-SARS-CoV-2 activity while chemically blocking MDR1 function, using the drug CP-100356 in the A549^+ACE2^ cell line and in HAEC (Fig. 7a). We observed no changes in antiviral efficacy when blocking MDR1 activity (Fig. 7b-d), suggesting that, in contrast to Vero E6 cells, this transporter does not play a role in our human airway model systems. Indeed, scRNA-seq analysis of HAECs (Fig. 5b^38^) did not reveal any detectable MDR1 (*ABCB1*) transcripts, suggesting levels of this transporter in HAEC are low.

**Figure 7.**
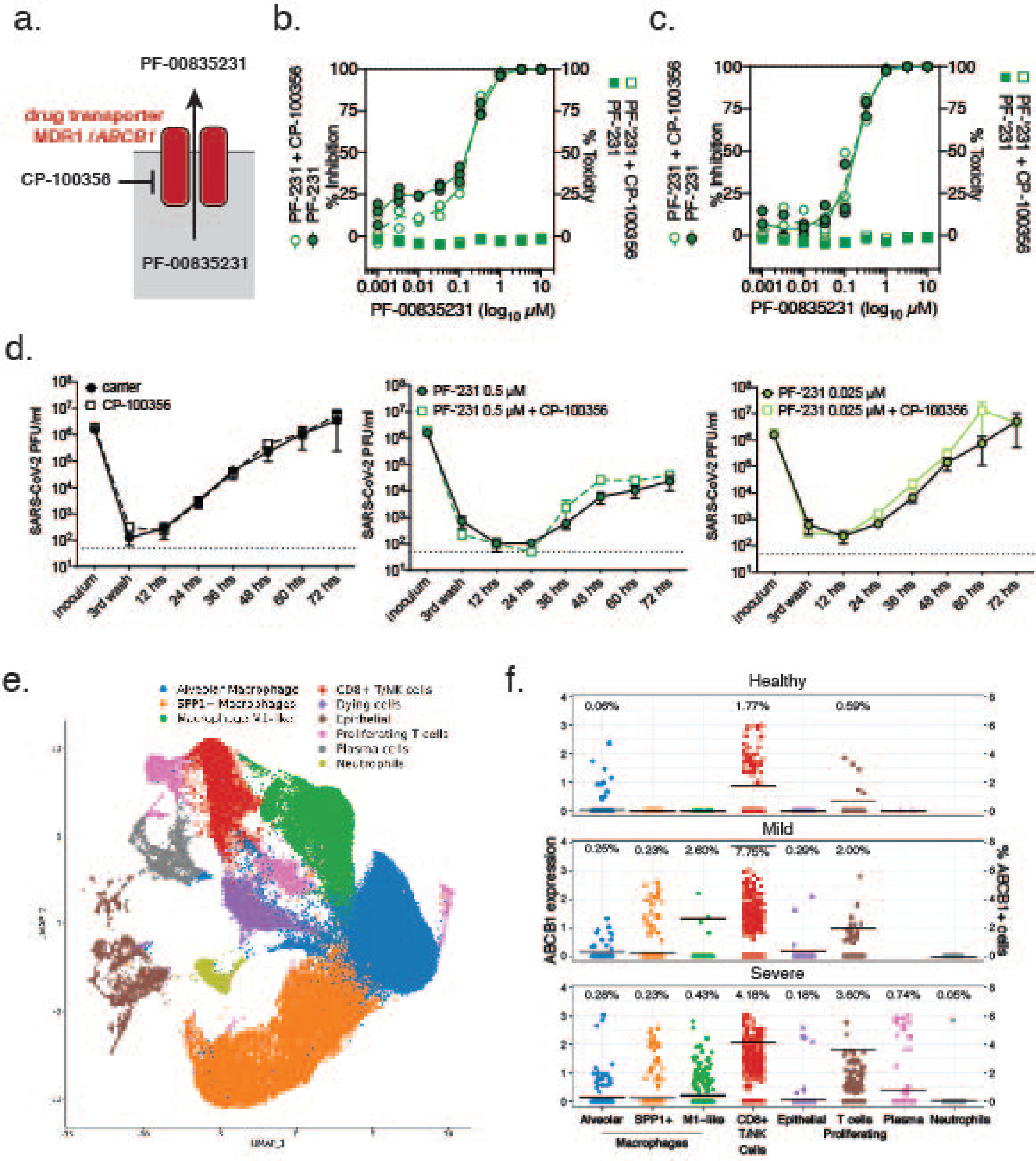
Role of MDR1 drug efflux transporter for PF-00835231-mediated SARS-CoV-2 inhibition in human airway cells. **a.** Schematic of experimental setup for (**b.-d.**). MDR1, encoded by ORF *ABCB1*, exports PF-00835231 from cells. CP-100356 was used as a chemical inhibitor to block MDR1 function. **b, c.** PF-00835231 antiviral activity and cytotoxicity in A549^+ACE2^ cells infected with SARS-CoV-2 USA-WA1/2020 for 24 (**b**.), or 48 hpi (**c.**) in the presence or absence of 1 µM MDR1 inhibitor CP-100356. Means ± SEM from n=3 independent experiments. **d.** Apical SARS-CoV-2 USA-WA1/2020 infectious titers from HAEC treated basolaterally with 0, 0.025, or 0.5 µM PF-00835231 in the presence or absence of 1 µM MDR1 inhibitor CP-100356. Means ± SEM from n=3 independent experiments. **e.** Clustered UMAP of single cells from bronchoalveolar lavages of n=12 patients. Integrated data of healthy patients (n=3) and COVID-19 patients with mild (n=3) or severe (n=6) symptoms. **f.** Normalized expression of *ABCB1* in clustered single cells from (**e.**). Left y-axis depicts level of ABCB1 expression; right y-axis depicts % of ABCB1-positive cells, also indicated by black bars; % of cells within each population with detectable ABCB1 transcripts shown above for each population.

A limitation of the previous experiments is that A549 cells originate from one patient, and HAEC were differentiated from precursor cells obtained from a single donor^41^. As a result, we cannot exclude that human genetic variation might influence MDR1 function or expression. Single nucleotide polymorphisms (SNPs) in the *ABCB1* promoter or within the open reading frame are well-described in the literature^15^. Furthermore, an inherent limitation of sc-RNAseq is that only abundant transcripts are detected, and given limited cellular heterogeneity of HAEC, it is possible that *ABCB1* transcripts remained undetected due to the limited depth of sequencing at single cell level.

To address these issues, we investigated transcript levels of *ABCB1* in bronchoalveolar lavages (BAL) of healthy individuals and of COVID-19 patients with mild or severe symptoms from a previously published dataset^42, 43^. BAL contained multiple cell types, including airway epithelial cells, but also immune cells, such as macrophages, T-cells, plasma cells, and neutrophils (Fig. 7e). In healthy individuals, we detected MDR1 (*ABCB1*) in 1.77 % of CD8+/NK cells, 0.59% of proliferating T-cells, and 0.06% of alveolar macrophages, but not in other cell types. Notably, we did not observe any *ABCB1* transcript in airway epithelial cells, which are thought to be the major site of SARS-CoV-2 replication^17, 18, 44, 45^. Compared to BAL from healthy individuals, BAL from COVID-19 patients with mild or severe symptoms showed an increased number of cells with detectable levels of MDR1 (*ABCB1*) transcripts (Fig. 7f). Interestingly, rather than an overall increase in *ABCB1* gene expression, this upregulation was limited to a small subset of individual cells, with the majority of cells remaining *ABCB1*-negative. This phenomenon was observed in alveolar macrophages (0.25%/0.28% positive in mild/severe COVID-19 patients), SPP1+ macrophages (0.23%/0.23%), M1-like macrophages (2.60%/0.43%), CD8+/NK cells (7.75%/4.18%), proliferating T-cells (2.00%/3.60%), plasma cells (cells not detected/0.74%), neutrophils (0%/0.05%) and in epithelial cells (0.29%/0.18%). Our findings suggest that although MDR1 (*ABCB1*) is upregulated in some cells during inflammatory processes such as observed in COVID-19, its expression remains cell type-specific, and only a small fraction of airway epithelial cells, the main replication sites for SARS-CoV-2, exhibit detectable levels of this efflux transporter.

Thus, we conclude that MDR1 is unlikely to significantly impact PF-00835231 efficacy during SARS-CoV-2 infection of the respiratory epithelium. In addition, our findings highlight the importance of using appropriate *in vitro* models for the evaluation of antiviral drugs.

## Discussion

The current public health emergency caused by COVID-19 has illustrated our dire need for vaccines and therapeutics to combat SARS-CoV-2. The SARS-CoV-2 polymerase complex is the target of the majority of small molecule inhibitors in multiple stages of development, including remdesivir^31^, favipiravir^20^, and ß-d-N4-hydroxycytidine^46^. At the time of writing, remdesivir is the only antiviral drug authorized for the treatment of COVID-19. In contrast to the abovementioned compounds, PF-00835231 blocks the SARS-CoV-2 3CL^pro^ protease^9^. PF-00835231 is the active component of PF-07304814, a first-in-class SARS-CoV-2 3CL^pro^ inhibitor currently in clinical trials. Here, we report the potent antiviral activity of PF-00835231 against SARS-CoV-2 in human lung epithelial cells and a model of polarized human airway epithelial cultures (HAEC). In our A549^+ACE2^ cell assay, we show that PF-00835231 has at least similar or better potency than the pre-clinical 3CL^pro^ inhibitor GC-376, or remdesivir. In HAEC, we find both remdesivir and PF-00835231 similarly potent.

The lack of inhibitors specific to SARS-CoV-2 early in the pandemic prompted off-label testing of protease inhibitors approved for other viruses, albeit with limited success^47^. This failure highlighted the need for novel compounds of greater specificity. A number of 3CL^pro^ inhibitors have since been identified and characterized in *in vitro* assays, including the cancer drug carmofur (1-hexylcarbamoyl-5-fluorouracil)^12^, an alpha-ketoamide inhibitor named 13b^7^, and a dipeptide-based inhibitor named GC-376^48^. GC-376, licensed for veterinary use^29^, was recently shown to inhibit SARS-CoV-2 in Vero E6 cells at an EC_50_ of 0.9 µM^11^. A different study showed PF-00835231 to inhibit SARS-CoV-2 at an EC_50_ of 0.27 µM in Vero E6 cells^9^. As such a comparison of historical data is problematic, we directly compared the antiviral efficacy of PF-00835231 and GC-376 side-by-side in the same assay (Fig. 3). We find that PF-00835231 is more potent that GC-376 in inhibiting SARS-CoV-2 and protecting cells from CPE. This result illustrates the *in vitro* potency of PF-00835231 compared to other, preclinical, 3CL^pro^ inhibitors, such as GC-376.

In coronaviruses, the genetic barrier to standard-of-care remdesivir or the pre-clinical drug ß-d-N4-hydroxycytidine is high, as mutations conferring resistance significantly reduce viral fitness, and cross-resistance between remdesivir or ß-d-N4-hydroxycytidine has not been documented^33, 46^. A high resistance barrier to 3CL^pro^-targeting drugs due to a high fitness cost has also been demonstrated for the beta-coronavirus murine hepatitis virus (MHV)^49^. Although the likelihood of selecting for resistant variants seems thus low, the existence of drugs with alternate targets may have important advantages. First, as seen in other acute and chronic viral infections, blocking multiple targets in combination therapy further decreases the likelihood for selection of viral resistance mutants^50–52^. Second, combination therapy with a cocktail of multiple drugs tackling different steps of the viral life cycle may have synergistic effects on controlling viral replication. Indeed, a recent *in vitro* study demonstrated significant synergistic effects between PF-00835231 and remdesivir in inhibiting SARS-CoV-2^14^. Third, upon failure of monotherapy, it is preferable to switch to an antiviral with a different target to circumvent cross-resistance caused by mutations in the same target^50–52^. For these reasons, the development of a diverse toolbox of antiviral drugs with different targets will be important to improve antiviral therapy in COVID-19.

The optimal window of opportunity for starting a successful antiviral drug regimen during acute viral infections, such as influenza, is the first few days post-symptom onset, while viral replication is actively ongoing^53^. For most COVID-19 patients, this window is likely limited to the first week of symptoms^54^. Such early treatment with remdesivir is impeded by its need for intravenous (IV) administration, requiring a healthcare facility setting, though it still demonstrated benefit for 68% of patients with more advanced infection in randomized clinical studies^55^. PF-07304814, with its active component PF-00835231 evaluated in this study, is also an IV treatment^9^. However, the time of active SARS-CoV-2 replication might be prolonged in the most severe patients, as suggested by the aforementioned clinical data of remdesivir^55^. This suggests the usefulness of SARS-CoV-2 antiviral regimens even at later times of infection, which further supports the investigation of PF-00835231 for the treatment of COVID-19.

P-glycoprotein (also known as MDR1, and encoded by gene *ABCB1*)^9^, is a membrane-associated ATP-dependent efflux pump capable of removing cytostatic drugs from target cells. The endogenous function of this transporter remains to be fully elucidated, but it is expressed across several immune cell types and other metabolically active cells. P-glycoprotein appears to be critical for maintenance and effector function of a range of cytotoxic immune cells^56–59^. Two recent studies suggested that PF-00835231 is a substrate for P-glycoprotein. This might pose a concern regarding the bioavailability of PF-00835231 in SARS-CoV-2-infected cells. We addressed this concern in two ways: functionally, by using chemical inhibition of MDR1 in human airway models, and transcriptionally, by determining the expression of *ABCB1* in HAEC or in cells obtained from BAL of healthy or COVID-19 patients. Our combined results demonstrate that MDR1 function does not impact PF-00835231 efficacy in our model systems, and is expressed at very low levels in airway epithelial cells, which are the major sites of SARS-CoV-2 replication^17, 18, 44, 45^. By mining scRNA-seq data from BAL of COVID-19 patients and healthy controls (n=12) for *ABCB1* expression, we considered the genetic variability on the *ABCB1* locus beyond that present in our two model systems. Indeed, a number of SNPs have been shown to alter expression of *ABCB1* transcripts, either by changing transcription factor or micro-RNA binding sites upstream of the *ABCB1* open reading frame, or by enhancing mRNA transcription due to silent mutations within the *ABCB1* open reading frame^15^. Other, missense, mutations within the *ABCB1* open reading frame have been shown to alter the functionality of MDR1, i.e. by altering membrane localization or recycling of the protein, or by modifying substrate recognition sites^15, 60^. Although we do not have information on the *ABCB1* genotype of our mined patient samples, the detected low expression of *ABCB1* transcripts in the cells permissive for SARS-CoV-2, together with the functional data from our model system, instills confidence that PF-00835231 efficacy is not hampered by the action of MDR1.

Spillovers of zoonotic coronaviruses with high pathogenic potential into the human population are not isolated events, as repeatedly illustrated by the emergence of SARS-CoV in 2002, Middle East Respiratory Syndrome Coronavirus (MERS-CoV) in 2012, and now SARS-CoV-2 in 2019^61^. To prepare for future pandemics, the development of pan-coronavirus compounds is of strategic importance. This involves choosing viral targets that are highly conserved within the coronavirus family, such as the 3CL^pro^ protease^7^. Indeed, a recent study performing in vitro protease activity assays with PF-00835231 revealed potent inhibition across a panel of diverse coronavirus 3CL^pro^, including those of alpha-coronaviruses (NL63-CoV, PEDV, FIPV), beta-coronaviruses (HKU4-CoV, HKU5-CoV, HKU9-CoV, MHV-CoV, OC43-CoV, HKU1-CoV), and a gamma-coronavirus (IBV-CoV)^14^. Our study revealed that the two 3CL^pro^ inhibitors tested (PF-00835231 and GC-376) were similarly potent for different clades of SARS-CoV-2 in which 3CL^pro^ is 100 % conserved. Of note, 3CL^pro^ is also 100 % conserved in the SARS-CoV-2 “UK variant” B.1.1.7 and the “Brazil variant” B.1.1.28, whereas the “South African variant” B.1.352 carries amino acid substitution K90R^62–64^. However, the K90 residue is distant from the active center of the protease, and is not expected to influence 3CL^pro^ substrate specificity. These findings support the notion that PF-00835231 is an inhibitor with broad coronavirus activity, which may be of use for emerging SARS-CoV-2 variants and for other emerging coronaviruses beyond SARS-CoV-2.

Together, our data from two human in vitro model systems for SARS-CoV-2 show efficient PF-00835231 antiviral activity and mitigate concerns arising from non-human models such as Vero E6 cells regarding PF-00835231 counteraction by the efflux transporter P-glycoprotein. Our results therefore inform and reinforce the ongoing clinical studies of pro-drug PF-07304814 and its active form PF-00835231 as a potential new treatment for COVID-19.

## Methods

### Study design

The primary goal of this study was to compare the *in vitro* efficacy and cytotoxicity of PF-00835231 and remdesivir in two human model systems for SARS-CoV-2 infection, A549^+ACE2^ cells and polarized human airway epithelial cultures. Compound characterization at NYU was done in a blinded manner. If not stated otherwise, all assays were performed in n=3 biological replicates. First, we performed in-depth characterization of A549^+ACE2^ cells for the study of SARS-CoV-2, using RT-qPCR, western blotting, flow cytometry, microscopy, and high-content imaging. Second, we evaluated the *in vitro* efficacy and cytotoxicity of PF-00835231, of a second, pre-clinical, protease inhibitor, GC-376, and of remdesivir in A549^+ACE2^ cells. We performed antiviral assays with SARS-CoV-2 from the two major clades during the first year of the COVID-19 pandemic. Third, we performed time-of-drug-addition assays in A549^+ACE2^ cells to delineate the time of antiviral action for PF-00835231, remdesivir and GC-376 within the SARS-CoV-2 life cycle. Fourth, we assessed the *in vitro* efficacy and cytotoxicity of PF-00835231 and remdesivir in the physiologically relevant model of polarized human airway epithelial cultures. Finally, we determined the role of efflux transporter MDR1 on the antiviral efficacy of PF-00835231. Our studies were intended to generate the data required to assess further pre-clinical investigations and the launch of a phase 1b clinical trial with PF-00835231 as a base-compound for the potential treatment of COVID-19.

### Cells and viruses

A549 cells were purchased from ATCC (cat no. CCL-185). To generate A549^+ACE2^ cells, we cloned the human ACE2 cDNA sequence (NP_001358344.1) into a pLV-EF1a-IRES-Puro backbone vector (Addgene, cat no. 85132), and prepared lentiviral particles as described previously^65^. A549 cells were transduced with pLV-EF1α-hACE2-IRES-Puro lentivirus and bulk-selected for transduced cells using 2.5 µg/mL puromycin. A549^+ACE2^ cells were maintained in DMEM (Gibco, cat no. 11965-092) containing 10% FBS (Atlanta Biologicals, cat no. S11150) (complete media), and puromycin (2.5 µg/mL final) was added to the media at every other passage. A549^+ACE2^ cells were used for SARS-CoV-2 infection studies. Vero E6 cells, purchased from ATCC (cat no. CLR-1586), were maintained in DMEM (Gibco, cat no. 11965-092) containing 10% FBS (Atlanta Biologicals, cat no. S11150), and were used for growing SARS-CoV-2 stocks and for SARS-CoV-2 plaque assays. Basal-like human airway progenitor cells (Bci-NS1.1^41^) were obtained from Dr. Ronald G. Crystal and were used for cytotoxicity assays and for the generation of polarized human airway epithelial cultures (HAEC). They were maintained in BEGM Medium (Lonza, cat no. CC-3171 and CC-4175) for cytotoxicity assays, while Pneumacult Ex Plus medium (StemCell, cat no. 05040) was used to culture Bci-NS1.1 cells for the generation of human airway epithelial cultures.

All SARS-CoV-2 stock preparations and following infection assays were performed in the CDC/USDA-approved BSL-3 facility in compliance with NYU Grossman School of Medicine guidelines for biosafety level 3. SARS-CoV-2 isolate USA-WA1/2020, deposited by the Center for Disease Control and Prevention, was obtained through BEI Resources, NIAID, NIH (cat no. NR-52281, GenBank accession no. MT233526). The USA-WA1/2020 stock, obtained at passage 4, was passaged once in Vero E6 cells to generate a passage 5 working stock (1.7E + 06 PFU/mL) for our studies on A549^+ACE2^. For studies on human airway epithelial cultures, passage 5 USA-WA1/2020 was amplified once more in Vero E6 cells and concentrated using an Amicon Ultra-15 centrifugal filter unit with a cut off of 100 kDa, resulting in a passage 6 working stock of 1.08E + 07 PFU/mL. SARS-CoV-2 USA/NYU-VC-003/2020 was isolated from patient nasal swab material in March 2020 (GenBank accession no. MT703677). We inoculated Vero E6 cells with a 1:2 dilution series of the nasal swab material in infection media (DMEM 2% FBS, 1% Pen/Strep, 1% NEAA, 10mM HEPES) to obtain passage 0 (P0) stock. P0 was passaged twice in Vero E6 to generate a passage 2 working stock (1.1E + 07 PFU/mL) for drug efficacy studies on A549^+ACE2^. For viral growth kinetics, pooled media from P0 stock was used to plaque purify a single virus clone on Vero E6 cells in presence of 1µg/ml TPCK-Trypsin, to avoid virus adaptation to Vero E6 cells due to the lack of TMPRSS2 expression^66^. Purified plaques were sequenced to verify the signature clade B amino acid changes, S D614G and NSP12 P323L, before expanding in presence of TPCK-Trypsin to generate a passage 1 working stock (1.8 E + 06 PFU/mL).

### Characterization of A549^+ACE2^ cells

Confluent 6-well A549 and A549^+ACE2^ cells were washed with PBS and cells were detached with CellStripper dissociation reagent (Corning cat no. 25056CI). Cells were pelleted, washed with PBS and either i) lysed in LDS sample buffer (ThermoFisher cat no. NP0007) supplemented with reducing agent (ThermoFisher cat no. NP0004) and Western blots were performed to analyze levels of ACE2 (1:1,000, GeneTex cat no. GTX101395) with beta-actin (1:10,000, ThermoFisher cat no. MA5-15739) as the loading control and imaged using Li-Cor Odyssey CLx, or ii) incubated in FACS buffer (PBS, 5% FBS, 0.1% sodium azide, 1mM EDTA) for 30 min on ice followed by 1 hour incubation with AlexaFluor 647 conjugated anti-ACE2 (1:40, R&D Biosystems cat no.FABAF9332R) or isotype control (1:40, R&D Biosystems cat no. IC003R) and subsequent analysis on CytoFLEX flow cytometer. Surface ACE2 was visualized by staining A549 and A549^+ACE2^ cells at 4°C with anti-ACE2 (1:500, R&D Biosystems AF933) and AlexaFluor 647 secondary antibody and DAPI. Images were collected on the BZ-X810 (RRID:SCR_016979, Keyence, Osaka, Japan) fluorescence microscope.

Confluent 6-well A549 and A549^+ACE2^ cells were collected in RLT lysis buffer supplemented with beta-mercaptoethanol and total RNA was extracted using Qiagen RNeasy mini kit. cDNA synthesis was performed using SuperScript™ III system (ThermoFisher cat no. 18080051) followed by RT-qPCR with PowerUp SYBR Master Mix (ThermoFisher cat no. A25742) on a QuantStudio 3 Real Time PCR System using gene-specific primers pairs for ACE2 and RPS11 as the reference gene. (ACE2fwd:GGGATCAGAGATCGGAAGAAGAAA, ACE2rev:AGGAGGTCTGAACATCATCAGTG, RPS11fwd:GCCGAGACTATCTGCACTAC, RPS11rev:ATGTCCAGCCTCAGAACTTC).

A549 and A549^+ACE2^ cells were seeded in black wall 96-well plates and at confluency, cells were infected with SARS-CoV-2. At 24 and 48 hpi, samples were fixed, stained with mouse monoclonal SARS-CoV anti-N antibody 1C7, which cross reacts with SARS-CoV-2 N (1:1000, kind gift of Thomas Moran), AlexaFluor 647 secondary antibody and DAPI, and imaged using CellInsight CX7 LZR high-content screening platform. Images were analyzed and quantified with HCS Navigator software. Syncytia were imaged using the Keyence BZ-X810 microscope at 60X magnification on A549^+ACE2^ cultured on chambered slides followed by 48 hpi SARS-CoV-2 infection and staining with SARS-CoV-2 N, AlexaFluor 647 secondary antibody, and DAPI.

### SARS-CoV-2 growth kinetics on A549^+ACE2^ cells

A549^+ACE2^ cells were seeded into 6-cm dishes at 70% confluency. The next day, media was removed and cells were washed twice with PBS with calcium and magnesium to remove residual medium. Cells were then infected at 0.01 multiplicity of infection (MOI), based on A549^+ACE2^ titer, at 37°C. The remaining inoculum was stored at −80°C for back titration. 1 hour post virus addition, virus was removed, cells were washed twice with PBS with calcium and magnesium to remove unbound virus and infection media (DMEM 2% FBS, 1% Pen/Strep, 1% NEAA, 10mM HEPES) was added. 60 μl of supernatant were collected and stored at −80°C to determine successful removal of input virus. Supernatant was then collected at 12, 24, 48 and 72 hpi, and stored at −80°C.

Viral titers in the supernatants were determined by focus forming assay. A549^+ACE2^ cells were seeded into black wall 96-well plates at 70% confluency. The next day, cells were then infected with 1:10 serial dilutions of the collected samples for 1 hour at 37°C. 1 hour post virus addition, virus was removed, and cells were overlayed with MEM 1.8% Avicell, 1% Pen/Strep, 1% GlutaMax, 20mM HEPES, 0.4% BSA, 0.24% NaHCO_3_. At 48 hours post infection, the overlay was removed, and cells were fixed by submerging in 10% formalin solution for 30-45 min. After fixation, cells were washed once with H_2_O to remove excess formalin. Plates were dried and PBS was added per well before exiting the BSL-3 facility. Fixed cells were permeabilized with Triton-X and stained with mouse monoclonal SARS-CoV anti-N antibody 1C7, which cross-reacts with SARS-CoV-2 N (kind gift of Thomas Moran), goat anti-mouse AlexaFluor 647, and DAPI. Plates were scanned on the CellInsight CX7 LZR high-content screening platform. A total of 9 images were collected at 4X magnification to span the entire well. Infection foci were counted manually.

### Human airway epithelial cultures (HAEC)

To generate HAEC, Bci-NS1.1 were plated (7.5 E + 04 cells/well) on rat-tail collagen type 1-coated permeable transwell membrane supports (6.5 mm; Corning, cat no. 3470), and immersed apically and basolaterally in Pneumacult Ex Plus medium (StemCell, cat no. 05040). Upon reaching confluency, medium was removed from the apical side (“airlift”), and medium in the basolateral chamber was changed to Pneumacult ALI maintenance medium (StemCell, cat no. 05001). Medium in the basolateral chamber was exchanged with fresh Pneumacult ALI maintenance medium every 2-3 days for 12-15 days to form differentiated, polarized cultures that resemble *in vivo* pseudostratified mucociliary epithelium. Cultures were used within 4-6 weeks of differentiation. HAEC were used for cytotoxicity assays and SARS-CoV-2 infections.

### Compound acquisition, dilution, and preparation

PF-00835231, remdesivir, and CP-100356 were solubilized in 100% DMSO and provided by Pfizer, Inc. Compound stocks diluted in DMSO to 30 mM were stored at −20°C. Compounds were diluted to 10 µM working concentration in complete media or Pneumacult ALI maintenance medium. All subsequent compound dilutions were performed in according media containing DMSO equivalent to 10 µM compound. GC-376 was purchased from BPS Biosciences (cat no. 78013) and used at 10 µM working concentration. SARS-CoV-2 (2019-nCov) rabbit polyclonal spike neutralizing antibody from Sino Biological (cat no. 40592-R001) was used at 3 µM working concentration. As a positive control for cytotoxicity assays, staurosporine was purchased from Sigma (cat no. S6942), and used at 1 µM working concentration.

### In vitro drug efficacy and cytotoxicity in A549^+ACE2^ cells

A549^+ACE2^ cells were seeded into black wall 96-well plates at 70% confluency. The next day, media was removed and replaced with complete media containing compound/carrier two hours prior to infection. Cells were then infected at 0.425 multiplicity of infection (MOI), based on Vero E6 titer, at 37°C. 1 hour post virus addition, virus was removed, and media containing compound/carrier was added. At 24 and 48 hours post infection, cells were fixed by submerging in 10% formalin solution for 30-45 min. After fixation, cells were washed once with H_2_O to remove excess formalin. Plates were dried and PBS was added per well before exiting the BSL-3 facility. Fixed cells were permeabilized with Triton-X and stained with mouse monoclonal SARS-CoV anti-N antibody 1C7, which cross-reacts with SARS-CoV-2 N (kind gift of Thomas Moran), goat anti-mouse AlexaFluor 647, and DAPI. Plates were scanned on the CellInsight CX7 LZR high-content screening platform. A total of 9 images were collected at 4X magnification to span the entire well. Images were analyzed using HCS Navigator to obtain total number of cells/well (DAPI stained cells) and percentage of SARS-CoV-2 infected cells (AlexaFluor 647 positive cells). To enable accurate quantification, exposure times for each channel were adjusted to 25% of saturation and cells at the edge of each image were excluded in the analysis. SARS-CoV-2-infected cells were gated to include cells with an average fluorescence intensity greater than 3 standard deviations that of mock infected and carrier treated cells. Representative images of viral foci were acquired using the BZ-X810 at 40X magnification of plates fixed at 48 hpi SARS-CoV-2 infection.

For determination of cytotoxicity, A549^+ACE2^ cells were seeded into opaque white wall 96-well plates. The following day, media was removed, replaced with media containing compound/carrier or staurosporine, and incubated for 24 or 48 hours, respectively. At these timepoints, ATP levels were determined by CellTiter-Glo 2.0 (Promega, cat no. G9242) using a BioTek Synergy HTX multi-mode reader.

### Time-of-drug-addition experiments

A549^+ACE2^ cells seeded into black wall 96-well plates and at confluency were treated and infected as followed. At 2.5 hours prior infection, cells were pre-treated with complete media containing 1x compound/carrier. In addition, SARS-CoV-2 (2x) was incubated with SARS-CoV-2 (2019-nCov) rabbit polyclonal spike neutralizing antibody (nAB, 2x). Pre-treated cells and virus/neutralizing antibody mix (1x) were incubated for 1 hour at 37°C. To synchronize infection, pre-incubated plates and SARS-CoV-2/nAB mix were chilled at 4°C for 30 min and SARS-CoV-2 was diluted on ice in media containing compound/carrier/nAB. Following pre-chilling, virus/compound/carrier/nAB mixtures were added to the cells to allow binding of virus for 1 hour at 4°C. Plates were moved to 37°C to induce virus entry and therefore infection. 1 hour post virus addition, virus was removed, and complete media was added to all wells. Complete media containing 2x compound/carrier/nAB was added to pre-treated cells, cells treated at infection and cells treated at 1 hour post infection. At 2, 3 and 4 hours post infection, complete media containing compound/carrier/nAb was added to according wells. At 12 hours post infection, samples were fixed, stained with SARS-CoV-2 N, AlexaFluor 647 secondary antibody, and DAPI, and imaged using CellInsight CX7 LZR high-content screening platform. Images were analyzed and quantified with HCS Navigator software as described for in vitro efficacy in A549+ACE2.

### In vitro efficacy and cytotoxicity in human airway epithelial cultures (HAEC)

48 hours prior to infection, 2-6-week-old HAEC were washed apically twice for 30 min each with pre-warmed PBS containing calcium and magnesium, to remove mucus on the apical surface. 2 hours prior to infection, HAEC were pretreated by exchanging the ALI maintenance medium in the basal chamber with fresh medium containing compounds or carrier. Remdesivir and PF-00835231 were used at 10, 0.5 and 0.025 µM, and CP-100356 at 1 µM. 1 hour prior to infection, cultures were washed apically twice for 30 min each with pre-warmed PBS containing calcium and magnesium. Each culture was infected with 1.35E + 05 PFU (Vero E6) per culture for 2 hours at 37°C. A sample of the inoculum was kept and stored at −80°C for back-titration by plaque assay on Vero E6 cells. For assessment of compound toxicity, additional cultures were washed and pre-treated as the infected cultures. Instead of being infected, these cultures were incubated with PBS containing calcium and magnesium only as Mock treatment. HAEC were incubated with the viral dilution or Mock treatment for 2 hours at 37°C. The inoculum was removed and the cultures were washed three times with pre-warmed PBS containing calcium and magnesium. For each washing step, buffer was added to the apical surface and cultures were incubated at 37°C for 30 min before the buffer was removed. The third wash was collected and stored at −80°C for titration by plaque assay on Vero E6 cells. Infected cultures were incubated for a total of 72 hours at 37°C. Infectious progeny virus was collected every 12 hours by adding 60 µL of pre-warmed PBS containing calcium and magnesium, incubation at 37°C for 30 min, and collection of the apical wash to store at −80°C until titration. Additionally, trans-epithelial electrical resistance (TEER) was measured in uninfected but treated HAEC to quantify the tissue integrity in response to treatment with compounds or carrier. At the end point, cultures were fixed by submerging in 10% formalin solution for 24 hours and washed three times with PBS containing calcium and magnesium before further processing for histology. Alternatively, at the end point, transwell membranes were excised and submerged in RLT buffer to extract RNA using the RNAeasy kit (Qiagen, cat no. 74104). cDNA synthesis was performed using SuperScript™ III system (ThermoFisher cat no. 18080051) followed by RT-qPCR with TaqMan universal PCR master mix (ThermoFisher cat no. 4305719) and TaqMan gene expression assay probes (ThermoFisher GAPDH cat no. 4333764F, BAX cat no. Hs00180269_m1, BCL2 cat no. Hs00608023_m1) using a QuantStudio 3 Real Time PCR System.

For additional determination of cytotoxicity in undifferentiated HAEC precursor cells, Bci-NS1.1 cells were seeded into opaque white wall 96-well plates. The following day, media was removed, replaced with media containing compound/carrier or staurosporine, and incubated for 24 or 48 hours, respectively. At these timepoints, ATP levels were determined by CellTiter-Glo 2.0 (Promega, cat no. G9242) using a BioTek Synergy HTX multi-mode reader.

### Histology on human airway epithelial cultures

For histology, transwell inserts were prepared using a Leica Peloris II automated tissue processor, paraffin embedded, and sectioned at 3 µm. The resulting slides were stained using a modified Periodic Acid–Schiff (PAS)-Alcian Blue protocol (Histotechnology,Freida L. Carson). Sections were imaged on the Leica SCN whole slide scanner and files were uploaded to the Slidepath Digital Image Hub database for viewing.

### Immunofluorescence on human airway epithelial cultures

For immunofluorescence of HAEC at top view, fixed and washed cultures were permeabilized with 50 mM NH_4_Cl (in PBS), 0.1% w/v saponin and 2% BSA (permeabilization/blocking (PB) buffer). Cultures were stained with i) rabbit polyclonal anti-SARS Nucleocapsid Protein antibody, which cross reacts with SARS-CoV-2 N (1:1000, Rockland cat no. 200-401-A50) and goat-anti-rabbit AlexaFluor 488, to visualize infection ii) mouse monoclonal anti-ZO-1-1A12 (1:500, Thermo Fisher cat no. 33-9100) and goat anti-mouse AlexaFluor 647 to visualize tight junctions, and DAPI. All dilutions were prepared in PB buffer. Images were collected on the Keyence BZ-X810 microscope.

### Single-cell RNA-seq analysis of human airway epithelial cultures

6-week-old human airway cultures were used for the single-cell RNA-seq analysis. The apical surface was washed once to remove mucus by adding 100 µL of PBS and incubating for 30 min at 37°C. Cells were dissociated by cutting out the transwell membrane and incubating it in 700 µL TrpLE 10x (Thermo Fisher cat no. A1217701) for 30 min rocking at 37°C. To increase dissociation cells were pipetted through wide-bore tips every 10 min during the incubation time. When cells were visually dissociated 700µl of ALI maintenance medium supplemented with 0.1% Pluronic (Thermo Fisher cat. No. 24040032) was added and cells were carefully pipetted again using wide-bore tips. The cell suspension was centrifuged through a 10 µM filter at 300 x g for 5 min to break up any remaining cell clumps. The cell pellet was washed once with 200 µL ALI maintenance medium supplemented with 0.1% Pluronic before cell number and viability was assed using the Countess II (Thermo Fisher) to calculate cell numbers used in the following steps for single-cell RNA-seq analysis.

Single-cell transcriptome profiling of dissociated organoids was carried out using the Chromium Next GEM Single Cell 5’ Library & Gel Bead Kit and Chromium controller (10X genomics). To enable multiplexing and doublet detection, cells were stained with barcoded antibodies described previously (Mimitou et al., 2019). Briefly, approximately 200,000 cells per sample were resuspended in staining buffer (PBS, 2% BSA, 0.01% Tween) and incubated for 10 minutes with Fc block (TruStain FcX, Biolegend; FcR blocking reagent, Miltenyi). Cells were then incubated with barcoded hashing antibodies for 30 min at 4 °C. After staining, cells were washed 3 times in staining buffer. After the final wash, cells were resuspended in PBS + 0.04% BSA, filtered, and counted. Cells were pooled and loaded onto the Chromium chips. For each lane, we pooled 5 samples, ∼10,000 cells per sample. The single-cell capturing, barcoding, and cDNA library preparation were performed using the Chromium Next GEM Single Cell 5’ Library & Gel Bead Kit by following the protocols recommended by the manufacturer. HTO additive oligonucleotide was spiked into the cDNA amplification PCR and the HTO library was prepared as described previously^67^.

The Cellranger software suite (https://support.10xgenomics.com/single-cell-gene-expression/software/pipelines/latest/what-is-cell-ranger) from 10X was used to demultiplex cellular barcodes, align reads to the human genome (GRCh38 ensemble, http://useast.ensembl.org/Homo_sapiens/Info/Index) and perform UMI counting. From filtered counts HTODemux was used to demultiplex hash-tagged samples and Seurat1 version 3.1.3 was used to process the single-cell data including dimension reduction, UMAP representation and differential expression to identify cell type specific markers determined by Wilcoxon test^68^.

### In silico analysis of bronchoalveolar lavages (BAL)

Filtered gene-barcode matrices for the BAL dataset were downloaded from GEO accession number GSE145926. Matrices were normalized using ‘LogNormalize’ methods in Seurat v.3 with default parameters and the resulting values were scaled using ScaleData. Seurat version 3.1.3 was used to process the single cell data including dimension reduction, UMAP representation and differential expression to identify cell type specific markers determined by Wilcoxon test.

### Statistical analysis

Antiviral activities of PF-00835231 and remdesivir in A549^+ACE2^ cells were determined by the following method. The percent inhibition at each concentration was calculated by ActivityBase (IDBS) based on the values for the no virus control wells and virus containing control wells on each assay plate. The concentration required for a 50% / 90% response (EC_50_ / EC_90_) was determined from these data using a 4-parameter logistic model. Curves were fit to a Hill slope of 3 when >3 and the top dose achieved ≥50% effect. Geometric means and 95% confidence intervals were generated in ActivityBase. Statistical comparisons were performed by log transforming the EC_50_ and EC_90_ values and fitting separate linear models to each endpoint, assuming equal log-scale variances across conditions and interactions of compound with strain and compound with time. The model can be described mathematically as

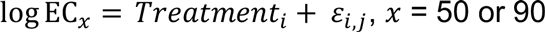

where *Treatment_i_* represents the effect of the combination of compound, strain, and time and *ε_i,j_* represents a normal error term for treatment *i* and assay replicate *j*. Contrasts between the factor combinations of interest were computed to assess significance and back-transformed into ratios of geometric means. Statistical significance was defined by a p value <0.05. Other statistical data analyses were performed in GraphPad Prism 7. Statistical significance for each endpoint was determined with specific statistical tests as indicated in each legend. For each test, a P-value < 0.05 was considered statistically significant.

## Data availability

All scRNA-seq data in this study can be accessed in GEO under the accession numbers GSE166601 and GSE145926.

## Acknowledgements

We would like to thank Thomas M Moran, Icahn School of Medicine at Mount Sinai, and Luis Martínez-Sobrido, Texas Biomedical Institute, for the kind gift of mouse monoclonal SARS-CoV N antibody 1C7. Histopathology of human airway cultures was performed by Mark Alu, Branka Brukner Dabovic and Cynthia Loomis from the NYUMC Experimental Pathology Research Laboratory. We are also grateful for to Adriana Heguy from the NYU Genome Technology Core. Both the Experimental Pathology Research Laboratory and the NYU Genome Technology Core are supported by NYU Cancer Center Support Grant P30CA016087, and NYU Langone’s Laura and Isaac Perlmutter Cancer Center. Statistical analysis of antiviral activities in A549^+ACE2^ cells was performed by Woodrow Burchett. We thank Ralf Duerr for critical reading of the manuscript. Research was further supported by grants from NIH/NIAID (R01AI143639 and R21AI139374 to MD, T32AI17647 to RAP, R01HL125816 to SBK), by Jan Vilcek/David Goldfarb Fellowship Endowment Funds to AMVJ, by The G. Harold and Leila Y. Mathers Charitable Foundation to MD, by a NYUCI Pilot Grant to SBK, by Pfizer Inc. to MD, and by NYU Grossman School of Medicine Startup funds to MD.

## Author contributions

MdV, ASM, JB, ASA, and MD conceived and designed the study. MdV, ASM, AMVJ, RAP, AS, PL, EI performed the experiments and analyzed the data. CS, RO, JB analyzed antiviral data. MdV, ASM, AMVJ, RAP, KR, SBK, LD, JB, MD interpreted the data. MdV, ASM, LD, SBK and MD wrote the paper.

## Materials and correspondence

All correspondence and material requests except those for antiviral compounds should be addressed to. Compound requests should be addressed to.

